# Incoherent merger network for robust ratiometric gene expression response

**DOI:** 10.1101/2022.11.07.515449

**Authors:** Ukjin Kwon, Hsin-Ho Huang, Jorge L. Chávez, Kathryn Beabout, Svetlana Harbaugh, Domitilla Del Vecchio

**Affiliations:** Department of Electrical Engineering and Computer Science, Massachusetts Institute of Technology, Cambridge, MA, USA; Department of Mechanical Engineering, Massachusetts Institute of Technology, Cambridge, MA, USA; 711th Human Performance Wing, Air Force Research Laboratory, Wright Patterson Air Force Base, OH, USA; UES, Inc., Dayton, OH, USA; Synthetic Biology Center, Massachusetts Institute of Technology, Cambridge, MA, USA

## Abstract

A ratiometric response gives an output that is proportional to the ratio between the magnitudes of two inputs. Ratio computation has been observed in nature and is also needed in the development of smart probiotics and organoids. Here, we achieve ratiometric gene expression response in bacteria *E. coli* with the incoherent merger network. In this network, one input molecule activates expression of the output protein while the other molecule activates an intermediate protein that enhances the output’s degradation. When degradation rate is first order and faster than dilution, the output responds linearly to the ratio between the input molecules’ levels over a wide range with *R*^2^ close to 1. Response sensitivity can be quantitatively tuned by varying the output’s translation rate. Furthermore, ratiometric responses are robust to global perturbations in cellular components that influence gene expression because such perturbations affect the output through an incoherent feedforward loop. This work demonstrates a new molecular signal processing mechanism for multiplexed sense-and-respond circuits that are robust to intra-cellular context.

## 1 Introduction

Ratiometric computation is common in biological systems since it allows cells to respond to relative, as opposed to absolute, changes in input stimuli (Fig. 1). For example, in the yeast sugar utilization network, the ratio between galactose and glucose controls the induction of galactose metabolic genes [1] [2]. In mammalian cells, the BMP signaling pathway comprises receptors that perform ratio sensing of multiple ligand concentrations, which ultimately leads to ratiometric responses of SMAD target genes [3] [4]. Finally, the ATP/ADP ratio drives the process of ATP hydrolysis and hence controls many reactions in the cell [5]. Ratiometric biomolecule signatures have also been linked to cognitive performance in humans [6] [7] [8] [9]. In particular, the ratio between norepinephrine and cortisol was shown to discriminate post-traumatic stress disorder (PTSD) patients from others [10] [11] and the ratio between cortisol and DHEAS is predictive of stress level [12] [13] [14]. For this reason, future smart probiotics could couple ratiometric biomarker sensors with genetic response systems that produce drugs to compensate for biomarker imbalances [15] [16][17] [18] [19]. Finally, in programmable organoids and directed cell differentiation, proper tissue composition requires the maintenance of specific ratios between different cell types’ population sizes, and hence the ability of computing and responding to ratios is required [20] [21].

**Figure 1:**
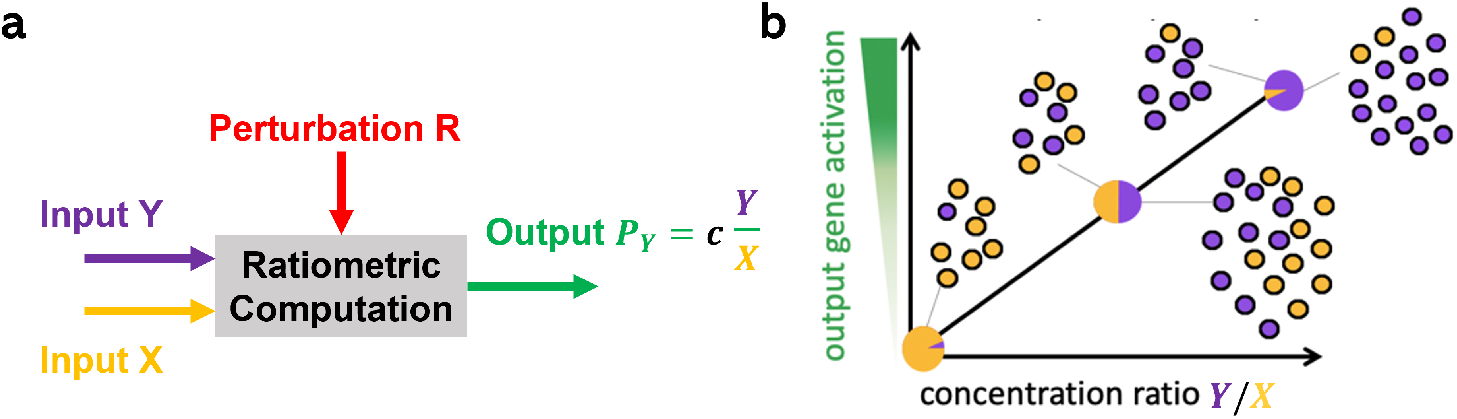
Robust ratiometric response. **(a)** A ratiometric response takes two signaling molecules as inputs (X and Y) and provides a protein P_*Y*_ as an output. The concentration of the output protein is proportional to the ratio between the concentrations of X and Y with proportionality constant *c*. R is a perturbation acting on the system that cannot be controlled and is unknown. If the output is independent of the perturbation R, the ratiometric response is robust to R. **(b)** Pictorial representation of the ratiometric input/output response. The output responds linearly to changes in the ratio *Y/X*.

Although natural ratiometric computation has been identified upstream of gene expression, notably through competitive receptor-ligand or promoter-regulator binding, it ultimately affects cellular functions through gene expression responses [1] [2] [3] [4]. These responses, in turn, have been shown to be robust to variations in the abundance of select cellular components in some natural networks [1]. On the other hand, gene expression is sensitive to variations in the levels of cellular resources, whose availability changes when genes become dynamically activated and repressed in the cell [22] [23] [24] [25]. Specifically, when a gene is activated, such as in response to one signaling molecule, any other gene’s expression rate, possibly responding to a different molecule, will decrease due to reduction of translation resources in bacteria and of transcription resources in mammalian cells [22][26]. As a result, although sensing through receptorligand or promoter-regulator binding may not be affected by changes in the availability of transcriptional and translational resources, the gene expression responses to these sensing events are influenced by them.

In this paper, we introduce the *incoherent merger network*, a design that performs ratiometric sensing on the gene expression output response directly. In this network, one of the two input molecules activates the expression of the output protein while the other activates an intermediate protein that enhances the degradation of the output. Because global variations in cellular resources implicated in gene expression equally affect both the output and intermediate protein, and the latter degrades the output, an incoherent feedforward loop “filters” the effect of such global perturbations on the output (Fig. 2(a)). As a consequence, the ratiometric output response can become robust to such global changes.

**Figure 2:**
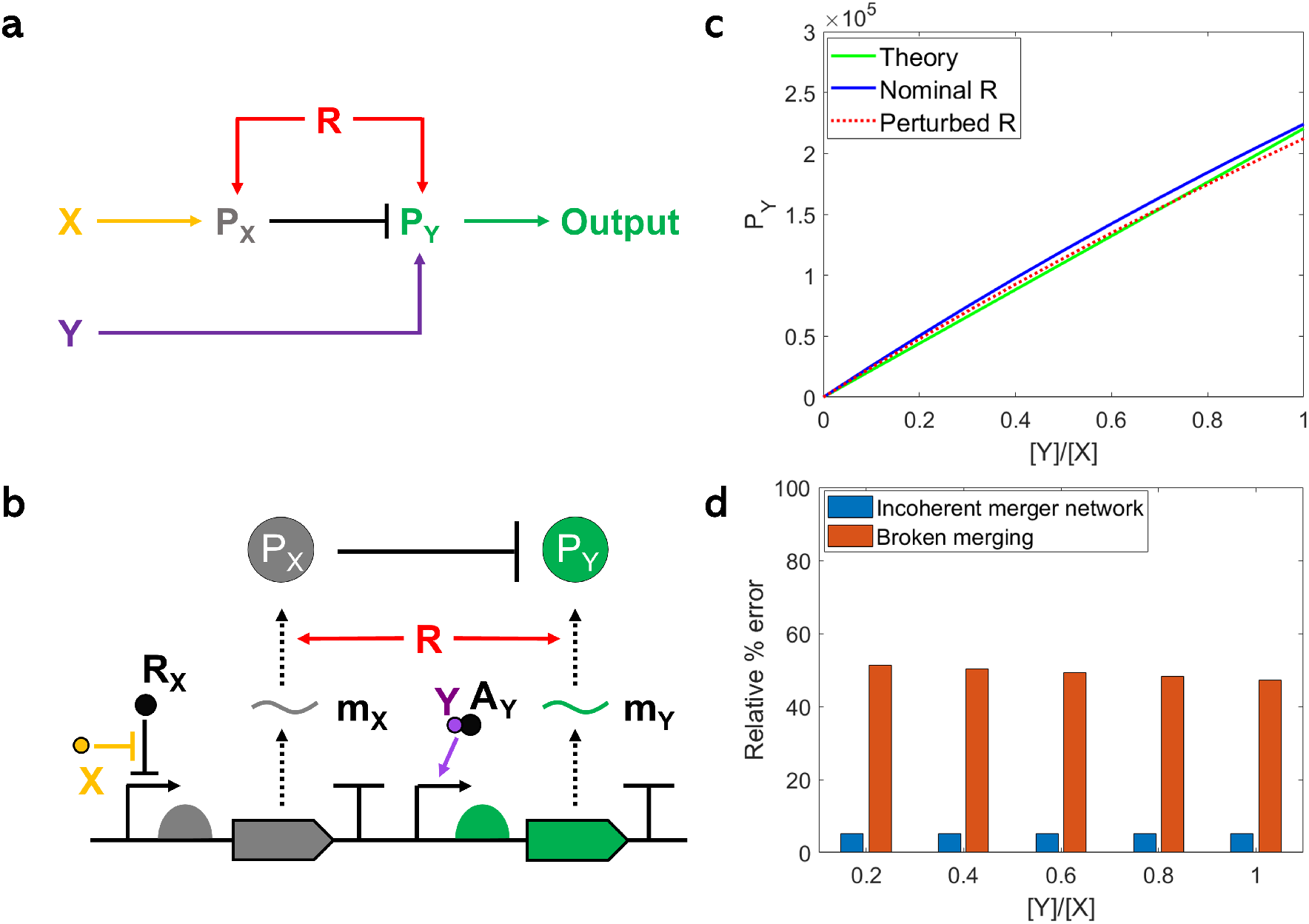
The incoherent merger network and its genetic circuit implementation. **(a)** Merger network motif with inputs X and Y and disturbance R, which affects both P_*X*_ and P_*Y*_ with the same sign. Here, arrow ‘→‘ denotes upregulation and ‘ ‘ represents downregulation. The loop formed by R, P_*X*_, and P_*Y*_ is an incoherent feedforward loop. **(b)** Genetic implementation of the network motif in **(a)**. Here, X is the first input, which is a negative inducer that binds to repressor protein R_*X*_ and prevents it from binding to DNA. Molecule m_*X*_ is the mRNA of P_*X*_. Signaling molecule Y is the second input, which is a positive inducer that binds to activator protein A_*Y*_ and allows DNA to be transcribed. Molecule m_*Y*_ is the mRNA of P_*Y*_. The half disks in front of the gene coding region represent RBS sequences. In panel (a), R is any cellular resource that is equally required for the expression of P_*X*_ and P_*Y*_, including transcriptional and translational resources. In the specific genetic implementation in (b), R is a translational resource, such as the ribosome. **(c)** Green colored plot is the theoretical steady state value of the output protein P_*Y*_ obtained from Supplementary Equation (28) with parameters in Supplementary Table 2 where assumptions (A0) - (A2) are satisfied. Blue and red colored plots are obtained from the detailed ODE model in Supplementary Equation (6) with parameters in Supplementary Table 2, in which perturbed *R* is 50% of nominal *R*. 50%change in *R* leads to only 5% change in the output, so the system attenuates the change. **(d)** Relative % error (=|*P*_*Y, Nominal R*_ − *P*_*Y, Perturbed R*_|*/P*_*Y, Nominal R*_ × 100 (%)) of incoherent merger network and broken merging, in which perturbed *R* is 50% of nominal *R*. In the broken merging, P_*X*_ does not degrade P_*Y*_.

We demonstrate this design with a bacterial genetic circuit where the intermediate molecule is a protease that targets the output protein for degradation. We develop a mathematical model of this system to determine the regime of biochemical parameters under which robust ratiometric response can be achieved. We experimentally demonstrate ratiometric computation and quantitative tuning of the response sensitivity based on model-driven part choices. We thus show that the sensitivity does not change when we vary the level of cellular resources through activation of a competitor gene. These results demonstrate a new mechanism to achieve ratio response, which is also robust to variability in global cellular components. More generally, our results can be used to engineer multiplexed genetic sense-and-respond systems that are robust to variable cellular context.

## 2 Results

### 2.1 The incoherent merger network requirements for robust ratiometric response

We first demonstrate the mechanism for robust ratiometric response by the incoherent merger network shown in Fig. 2(a) by using a simple ordinary differential equation (ODE) model that describes the rate of change of the species concentration. For any species S, we denote in italics *S* its concentration. Here, X is a signaling molecule that enables the production of protein P_*X*_ and this production requires the cellular resource R. Resource R can model any transcriptional or translational resource, such as RNAP or ribosomes, and we leave it unspecified at this time. Similarly, Y is a signaling molecule that enables the production of the output protein P_*Y*_, which also requires the same resource R for its production. Assuming that the production of P_*X*_ and of P_*Y*_ can be written as one-step enzymatic reactions, we have:

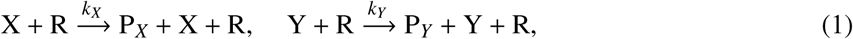

where *k*_*X*_ and *k*_*Y*_ are production rates constants. The interaction P_*X*_ ⊣ P_*Y*_ in Fig. 2(a) can be implemented by either having P_*X*_ enhance the degradation of P_*Y*_ or inhibit its production. Here, we choose the former and take P_*X*_ as a protease, which binds to its target on protein P_*Y*_, forming a complex C, which leads to the release of P_*X*_ and degradation of P_*Y*_ [27]:

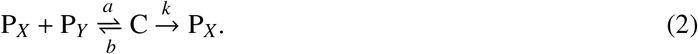

We also consider dilution, 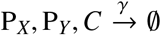 where *γ* is the dilution rate constant. By assuming the complex concentration *C* at the quasi-steady state and by letting *P*_*XT*_ = *P*_*X*_ + *C* be the total concentration of P_*X*_, we obtain the reduced ODE model:

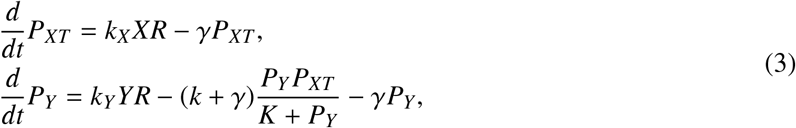

where *K* = (*b* + *k* + *γ*)*/a* is the Michaelis-Menten constant of the protease reaction (5). To achieve robust ratiometric sensing, we require that (A1) *P*_*Y*_ ≪ *K* and (A2) *γ* ≪ *P*_*XT*_ (*k* + *γ*)*/K*. With these assumptions, we obtain that the steady state value of the output protein P_*Y*_ is given by

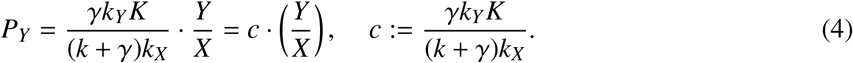

Therefore, the steady state value of the output protein P_*Y*_ is proportional to the ratio *Y/X* and this ratio is independent of the resource level *R*. In summary, the key design requirements to achieve robust ratiometricsensing are

(A0) The production rates of P_*X*_ and P_*Y*_ are linear with the levels of the signaling molecules X and Y, respectively, and with the resource level *R* (satisfied in this model by reactions (1));

(A1) The protease enzymatic reaction is in the first order regime, that is, *P*_*Y*_ ≪ *K*;

(A2) Dilution rate is negligible with respect to degradation rate, that is, *γ* ≪ *P*_*XT*_ (*k* + *γ*)*/K*.

Regarding (A0), the linearity of the production rates of P_*X*_ and P_*Y*_ with the levels of X and Y, respectively, is needed for ratiometric computation, while the linearity of the production rates with the level of cellular resource R is needed to make the level of P_*Y*_ independent of *R*. Assumptions (A1) and (A2) can be both satisfied by taking a sufficiently strong protease (large catalytic constant *k*). Finally, we can tune the sensitivity *c* by varying the production rate constant *k*_*Y*_ of protein P_*Y*_, which can be easily accomplished by varying the ribosome binding site (RBS) strength for P_*Y*_.

We consider the specific genetic implementation of the incoherent merger network shown in Fig. 2(b). In this implementation, X and Y are sensed through transcriptional sensors, wherein X and Y each bind to a transcriptional regulator to modulate transcription of the regulator’s target promoter. Transcriptional sensors have been developed for a variety of signaling molecules and therefore there is a large library to select from [28] [29] [30] [31] [32] [33]. Although we focus our analysis on transcriptional sensors, translational sensors are also an option [34] [35]. For the sake of illustration, we choose X and Y as a negative and as a positive inducer, respectively. Specifically, a negative inducer is a molecule that binds to a transcriptional repressor and prevents it from binding to DNA and hence to obstruct RNAP binding. A positive inducer, instead, is a small molecule that binds to a transcriptional activator and allows this activator to bind to DNA/RNAP to start transcription [27]. We also specialize resource R to a translational resource, such as ribosomes. In fact, it is well known that in bacterial synthetic genetic circuits translational resources are responsible for the observed interference among independently regulated genes [22] [23].

Our detailed mathematical model of the genetic circuit in Fig. 2(b) reveals parameter conditions under which assumption (A0) can be satisfied (Supplementary Equations (8)-(23)). Specifically, the linearity of the production rates of P_*X*_ and P_*Y*_ in X and Y, respectively, is approximately satisfied within a range of X and Y concentrations. This range can be widened by suitable choices of biochemical parameters of the transcriptional sensors, such as the regulators’ total concentration and target DNA copy number (Supplementary Equations (8)-(10), (13)-(15)), and can be empirically determined by inspecting the experimental dose response curves of the sensors as we do in Section 2.2. The linearity in *R* of the production rates of P_*X*_ and P_*Y*_ is ensured when both the repressor’s and activator’s mRNAs are saturated by ribosomes while the mRNAs of P_*X*_ and P_*Y*_ are not. Suitable choices of the RBS strengths achieve these requirements, that is, sufficiently strong RBS for the repressor and activator proteins and weaker for the output protein P_*Y*_ and protease P_*X*_ (Supplementary Equations (19)-(23)). Assumptions (A1) and (A2) can be satisfied by taking a sufficiently strong protease, which is equivalent to large catalytic constant *k*. The slope *c* can also be quantitatively tuned by varying the strength of the RBS of the output protein P_*Y*_ (Supplementary Equation (28)). Satisfying the assumptions (A0) - (A2) (Supplementary Table 2) confirm that ratiometric sensing is robust to changes in the level of resource R, and that the steady state value of the output protein P_*Y*_ from the detailed ODE model in Supplementary Equation (6) can be approximated as Supplementary Equation (28) (Fig. 2(c)). Also, degradation of P_*Y*_ by P_*X*_ is critical for the circuit’s output is robust to changes in the level of resource R (Fig. 2(d)). For both Figs. 2(c)-(d), R perturbed is 50% of R nominal, which is reasonable assumption according to [22].

### 2.2 The incoherent merger network achieves tunable ratio computation

We chose two transcriptional sensors from the Marionette strain [28] and a protein degradation tag from the library developed in [36]. For demonstration, we chose transcriptional sensors that satisfy (A0) in a large range of X and Y concentrations (Supplementary Figure 7a). In general, for a given transcriptional sensor, there will be an acceptable linear range of the signaling molecule level. We can make this range larger by suitably changing key biochemical parameters, such as the total regulator (activator or repressor) level and the copy number of the regulator DNA target (Supplementary Note 1.1). We thus chose IPTG as X and Sal as Y and found a linear range from 0 to 50 *µ*M for the Sal sensor and from 50 to 400 *µ*M for the IPTG sensor (Supplementary Figure 7b). As P_*X*_, we chose the *mf* -Lon protease and picked the pdt#3 tag, which has the strongest protein degradation as experimentally determined in [36]. Finally, we chose superfold GFP (sfGFP) as P_*Y*_ (Fig. 3(a)). We transformed the constructs into the host cell *E. coli* Marionette strain in which the required regulators, LacI (R_*X*_) and NahR (A_*Y*_), are chromosomally integrated [28].

**Figure 3:**
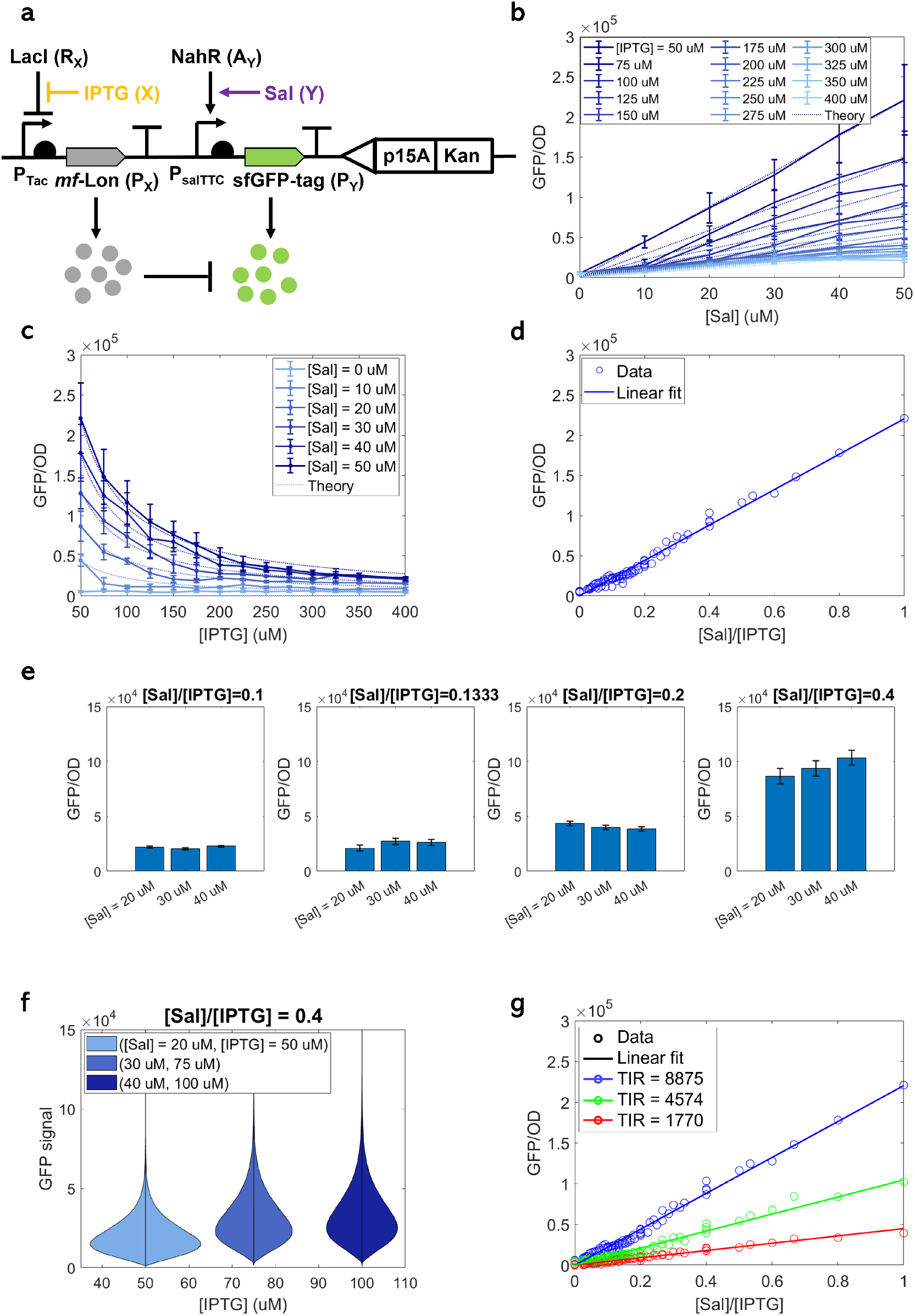
Performance and tunability of the incoherent merger network genetic implementation. **(a)** Genetic diagram of the incoherent merger network. This genetic circuit was created in three variants, depending on the choice of the RBS for sfGFP. Specifically, we chose TIRs as 1770, 4574, 8875 (Supplementary Table 1 and Supplementary Figure 1). Both LacI and NahR regulators are endogenously expressed from the host *E. Coli* Marionette strain [28]. **(b)** Dose-response curves showing how GFP depends on Sal along with theoretical model predictions from the model in Supplementary Equation (28) with parameter values in Supplementary Table 2. **(c)** Dose-response curves showing how GFP depends on IPTG along with theoretical model predictions from the model in Supplementary Equation (28) with parameter values in Supplementary Table 2. **(d)** Response to the signal ratio *Y/X*. GFP/OD value of each circle is an average of 3 biologically independent replicates. **(e)** Response to the signal ratio *Y/X* for selected [Sal] and [IPTG] combinations in **(d)** that have the same ratio. **(f)** Distributions of output levels per cell for selected [Sal] and [IPTG] combinations that have the same ration (0.4). Coefficient of variation (CV) is 0.5374, 0.5025, 0.5007 for [IPTG] = 50, 75, 100 *µ*M, respectively. All data were measured by flow cytometry when OD value was close to 0.054. Distributions of output levels per cell for all [IPTG] and [Sal] combinations are shown in Supplementary Figure 4 and Supplementary Figure 5. **(b)-(f)** were obtained with sfGFP’s TIR ± = 8875. **(g)** Tunability of the ratiometric sensor. Blue, green and red colored plots represent TIR = 8875, 4574, 1770, respectively. The temporal growth data and GFP expression with different TIRs can be found in Supplementary Figure 8 - Supplementary Figure 10, respectively. Data in the line plots in panel (b)-(c) and data in the scatter plot in panel (d) and (g) represent mean values (±SD) of n = 3 biologically independent experiments.

We characterized the incoherent merger network response in the indicated linear range of the respective IPTG (X) and Sal (Y) concentrations (Figs. 3(b) and (c)). The linearity between *P*_*Y*_ ([sfGFP]) and *Y* ([Sal]) is apparent from Fig. 3(b), and the required *P*_*Y*_ ∝ 1*/X* trend between *P*_*Y*_ and *X* ([IPTG]) is apparent from Fig. 3(c). In particular, for both induction curves, experimental results match well the theoretical model predictions. A scatter plot of *P*_*Y*_ versus the ratio *Y/X* shows a linear relationship with an *R*^2^ value of 0.98347, demonstrating the expected performance of the ratiometric sensor (Fig. 3(d)). To further demonstrate that the output responds to changes in the ratio but not to changes in *X* or *Y* alone when the ratio is constant, we selected different pairs of *X* and *Y* that have the same ratio and plotted bar charts of the corresponding outputs (Fig. 3(e)). These data confirm that the output remains approximately constant when the ratio of *X* and *Y* stays the same.

It is well known that ratio computation is highly sensitive to noise in *X* especially when *X* is small [37]. To determine how randomness due to intrinsic noise can affect ratiometric sensing, we conducted flow cytometry experiments (Supplementary Figure 4 and Supplementary Figure 5) and obtained scatter plots of *P*_*Y*_ versus the ratio *Y/X* (Supplementary Figure 6), which also shows a linear relationship between *P*_*Y*_ and *Y/X* with an *R*^2^ value of 0.925. Violin plots of the output GFP fluorescence for decreasing values of IPTG show that in the tested inducer concentration ranges, noise level remains contained. Specifically, values of the coefficient of variation (CV) remain approximately constant through the range of IPTG concentrations (Fig. 3(f)).

To show the tunability of the response, we created a library of three constructs, where the strength of the GFP RBS measured by the translation initiation rate (TIR) using the RBS Calculator [38] is progressively decreased from 8875, to 4575, to 1770 (Supplementary Table 1, Supplementary Figure 1 and Supplementary Figure 2). According to [38], the TIR is proportional to parameter 1*/K*_6_ = *a*_6_*/*(*b*_6_ + *k*_6_ + *γ*) and the slope *c* is proportional to 1*/K*_6_ (Supplementary Equation (12)). Therefore, we expect that the slope should progressively decrease from 220,428, observed in Fig. 3(d), to 104,594, to 44,846, which is validated by the experimental data (Fig. 3(g)). Specifically, when the TIR was changed from 8875 to 4574, the slope decreased from 220,428 to 104,594, which is a 47.5% decrease in the slope, consistent with a 51.5% decrease in the TIR. Similarly, when the TIR was changed from 8875 to 1770, the slope decreased from 220,428 to 44,846, which is a 20.3% decrease in the slope, consistent with a 19.9% decreaase in the TIR. This data indicate that the sensitivity of our ratiometric sensor is quantitatively tunable through modulation of the strength of the RBS of the output protein.

In order to demonstrate that ratio computation hinges on the incoherent merging of the X and Y signals onto P_*Y*_, we created a control construct in which the protease tag was removed from the GFP protein, thereby removing the enhanced degradation of GFP by the protease (Fig. 4(a) and Supplementary Figure 3). In this case, we still expect that the level of GFP decreases as X is increased, due to sequestration of ribosomes by P_*X*_ [22], but the model modifies to (see Supplementary Note 1.2):

**Figure 4:**
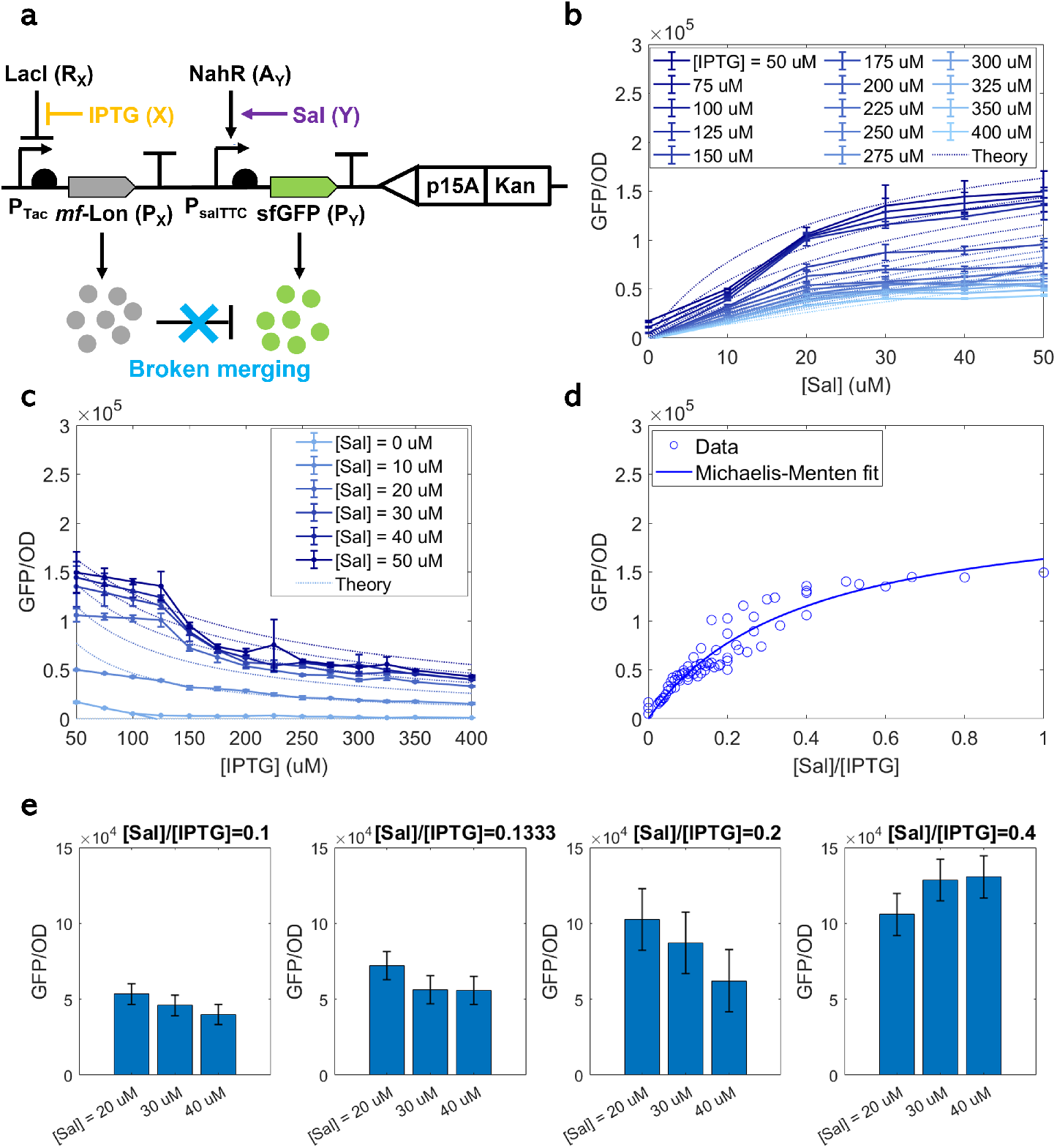
Breaking the merging breaks ratiometric computation. **(a)** Genetic diagram of the circuit where GFP misses the degradation tag. **(b)** Dose-response curves showing how GFP depends on Sal along with theoretical model predictions from the model in (30) with parameter values in Supplementary Table 2. **(c)** Dose-response curves showing how GFP depends on IPTG along with theoretical model predictions from the model in (30) with parameter values in Supplementary Table 2. **(d)** GFP response to the signal ratio *Y/X*. GFP/OD value of each circle is an average of 3 biologically independent replicates. **(e)** Response to the signal ratio *Y/X* for selected [Sal] and [IPTG] combinations in **(d)** that have the same ratio. **(b)-(e)** were obtained with sfGFP’s TIR = 8875. The temporal data of exponential growth and GFP expression can be found in Supplementary Figure 11. Data in the line plots in panel (b)-(c) and data in the scatter plot in panel (d) represent mean values (± SD) of n = 3 biologically independent experiments.

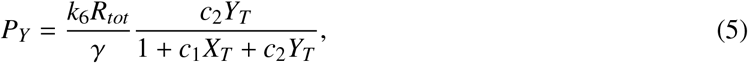

where *c*_1_ and *c*_2_ are defined in Supplementary Equation (30) and *R*_*tot*_ is the total concentration of ribosomes. This is not a ratio computation as confirmed by the experimental data of Fig. 4. Specifically, for small *Y, P*_*Y*_ linearly increases as *Y* increases, but it saturates for large *Y* (Fig. 4(b)). Dashed lines in Figs. 4(b) and (c) show theoretical model predictions from the model in (30) with parameter values in Supplementary Table 2. For both cases, experimental results match the theoretical predictions. Finally, different from Fig. 3(e), the output responds to changes in *X* or *Y* even if the ratio is constant since different pairs of *X* and *Y* that have the same ratio give different outputs (Fig. 4(e)). This confirms that, when the signal merging is broken, ratiometric computation is disrupted.

### 2.3 The incoherent merger network response is robust to perturbations in cellular context

Sophisticated genetic circuits typically include multiple sensors, logic computation/signal processing, and actuation [39] [40]. Because all of these components share cellular resources required for gene expression, independent circuit components can interfere with each other’s functionality [22] [23]. Here, we investigate to what extent the incoherent merger network response is robust to depletion of gene expression resources due to activation of other sensors or circuit modules in the cell. We achieve this by adding to the genetic system of Fig. 3(a) a resource competitor constituted of an inducible RFP gene (Fig. 5(a)). Referring to the incoherent merger network diagram of Fig. 2(a), reduced resource *R* decreases the production rate of both P_*X*_ and P_*Y*_. Since the concentration of P_*X*_ decreases, the degradation rate of P_*Y*_ also decreases. If the network is in the proper parameter regime, this compensation can completely cancel the effect of a decrease in *R* on *P*_*Y*_.

**Figure 5:**
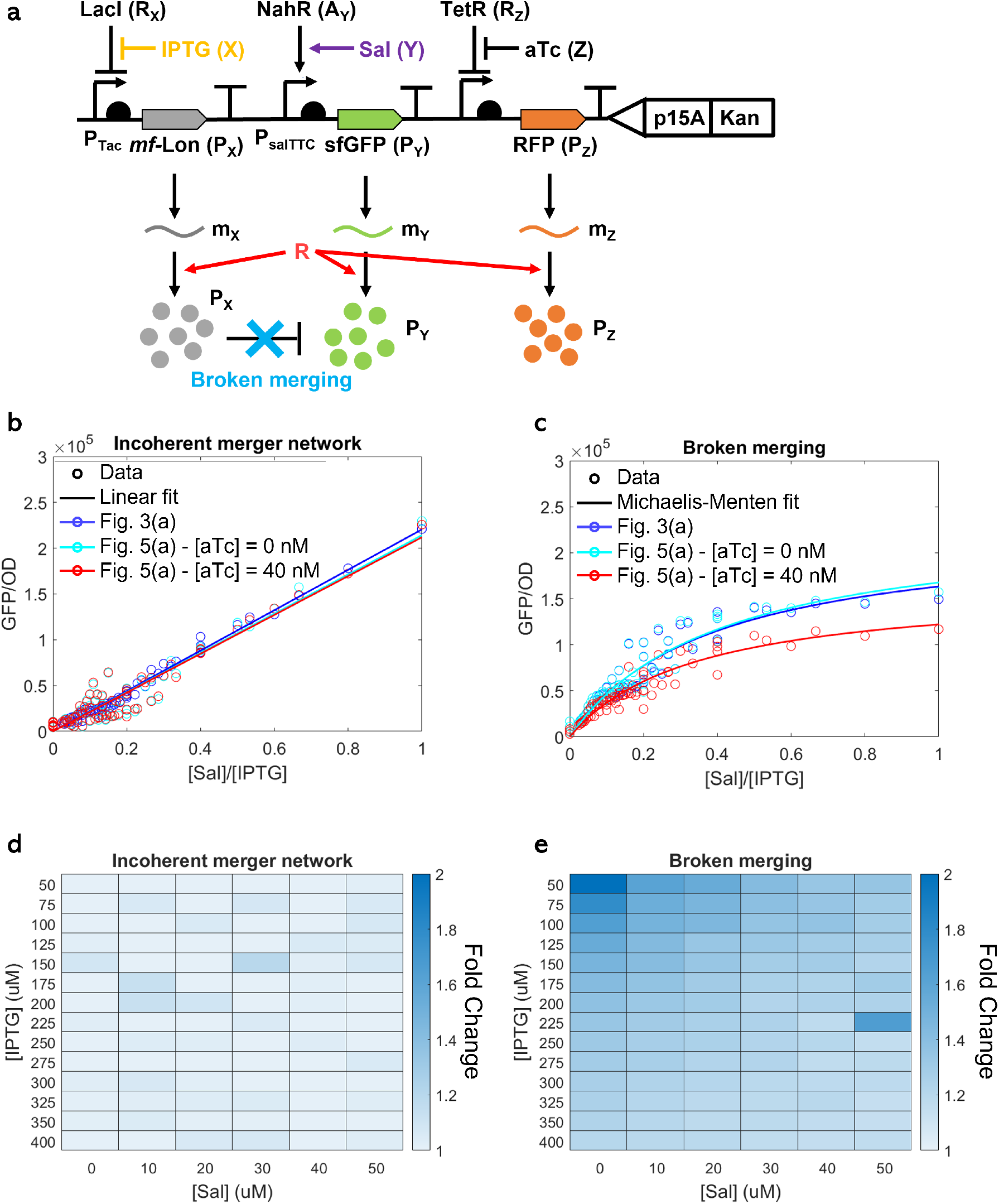
The incoherent merger network is robust to perturbations in translational resources. **(a)** Genetic diagram of the incoherent merger network with a resource competitor. “Broken merging” indicates that the GFP is devoid of the degradation tag and hence is not degraded by the protease. In the resource competitor, RFP expression is activated by addition of the aTc inducer. **(b)** Response of the incoherent merger network to the signal ratio *Y/X*. Blue colored plot shows the network without the resource competitor and cyan and red colored plots show the network with the resource competitor, with [aTc] = 0 nM and [aTc] = 40 nM, respectively. All solid lines represent linear fit. **(c)** Response of the broken merging to the signal ratio *Y/X*. Blue colored plot shows the network without the resource competitor and cyan and red colored plots show the network with the resource competitor, with [aTc] = 0 nM and [aTc] = 40 nM, respectively. All solid lines represent Michaelis-Menten fit. **(b)-(c)** GFP/OD value of each circle is an average of 3 biologically independent replicates. **(d)** Heat map showing the fold change of the steady state value of the output protein P_*Y*_ of the incoherent merger network when aTc is added (40 nM) with varying IPTG and Sal concentrations. **(e)** Heat map showing the fold change of the steady state value of the output protein P_*Y*_ of the broken merging when aTc is added (40 nM) with varying IPTG and Sal concentrations. **(b)-(e)** were obtained with sfGFP’s TIR = 8875. The temporal data of exponential growth and GFP expression of the incoherent merger network and the broken merging can be found in Supplementary Figure 12 and Supplementary Figure 13, and Supplementary Figure 14 and Supplementary Figure 15, respectively. Data in the scatter plot in panel (b)-(c) represent mean values (± SD) of n = 3 biologically independent experiments.

We conducted experiments to compare the steady state value of the output GFP protein of the circuit in Fig. 5(a) with either no competitor (RFP, [aTc] = 0 nM) or with high levels of competitor ([aTc] = 40 nM). The experimentally obtained plots of P_*Y*_ (GFP) versus the ratio *Y/X* ([Sal]/[IPTG]) show that the competitor and no competitor plots almost perfectly overlap (compare the cyan colored plot with the red colored plot). Both of these plots also overlap with the same plot obtained for the original circuit in Fig. 3(a) (blue colored plot), in which the RFP gene expression cassette is absent. By contrast, when the degradation tag on the GFP gene is removed (“broken merging” in Fig. 5(a)), not only ratiometric computation is lost, consistent with what observed in Fig. 4(d), but also the GFP output drops when the RFP gene is activated (compare the cyan and red colored plots in Fig. 5(c)). These data demonstrate that the presence of an incoherent feedforward loop encompassing the resource results in an output that is robust to changes in cellular resources and, hence, also in robust ratiomteric response. These data is also consistent with our model predictions, according to Supplementary Equations (28), (30) and (45).

To further show that this robustness to changes in cellular resources holds for every combination of the inputs X (IPTG) and Y (SAL), we also report a heat map in Figs. 5(d)-(e) showing the fold-change of the steady state of the output GFP level when RFP is expressed for the incoherent merger network and for the case when GFP has no degradation tag (broken merging). While for the incoherent merger network the foldchange never goes above 1.2, for the system with broken merging, the fold-change reaches 2 fold. These data show that ratiometric computation by the incoherent merger network is robust to changes in the level of cellular resources required for gene expression, which is not the case for the broken merging system.

## 3 Discussion

We have introduced the incoherent merger network to achieve robust ratiometric gene expression response. The output protein of the network responds to the ratio between the levels of two input molecules, but not to absolute changes that keep the ratio constant, and the response sensitivity can be quantitatively tuned (Fig. 3 and Fig. 4). The response is further robust to changes in global cellular components that affect gene expression, such as due to dynamic activation or repression of other genes (Fig. 5).

A well known molecular mechanisms for ratiometric sensing identified in natural systems is competitive binding of two input molecules to an integrator molecule, such as ligands binding to a receptor or regulators binding to a promoter [1][3]. This mechanism was leveraged to design a synthetic reporter that responds to the ratio ATP/ADP with a Michaelis-Menten function [5] and was later implemented with single guide RNAs (sgRNAs) for integrating the signals from two competing sgRNAs into one reporter output [41]. In these designs, the molecule that integrates the signals (a promoter) is different from the molecule that responds to the signal ratio (the output protein), therefore the output response will vary as the level of global cellular resources implicated in gene expression change. In the incoherent merger network design, instead, the molecule that integrates the signals is the gene expression output protein itself and integration occurs through protein-protein interaction via modulation of protein activity. This ensures that the output’s response is robust to any global perturbations in cellular components implicated in gene expression.

Although the incoherent merger network motif has not been linked before to ratiometric computation, it is found in all those incoherent feedforward loops (iFFL) of type I [42] where repression occurs through modulation of protein activity via protein-protein interaction. Many such iFFLs have been reported in natural networks and implicated in the control of cellular functions, such necroptotic cell death [43], initiation of mitotic exit [44], cell migration [45], and virulence in *Salmonella* [46]. In the necroptotic cell death iFFL, the output protein RIPK3 leads to plasma membrane rupture and cell death, and its activity is modulated by the A20 protein via protein-protein binding [43]. In turn, RIPK1 is an upstream regulator of RIPK3 and the A20 protein has many upstream regulators [47]. These A20 regulators and RIPK1 are inputs to the incoherent merger network and hence RIPK3 activity may respond to relative, but not to absolute, changes in the levels of these inputs. Similarly, in the iFFL implicated in cell cycle regulation, the output is the mitotic exit protein Dbf2, which initiates mitotic exit, and the Cdk1 kinase induces Dbf2 enhanced degradation by phosphorylation [44]. Cdk1’s activity is, in turn, modulated by the CDC25C phosphatase and Dbf2 has several upstream regulators, including Mob1 and CDC15 [48] [49]. Since the CDC25C phosphatase and Mob1/CDC15 enter the incoherent merger network as inputs, it is possible that Dbf2’s activity responds to relative changes between CDC25C and Mob1/CDC15 levels but not as much to absolute variations. Finally, in the iFFL controlling virulence in *Salmonella*, the HilD protein controls the fine balance between growth cost and virulence benefit and its activity is regulated by the HilE protein through protein-protein interaction [46]. Both HilD and HilE have many known regulators that enter the incoherent merger network as inputs. It is therefore possible that the balance between growth cost and virulence benefit is dictated by the relative concentrations among these regulators. Therefore, all the above systems could be re-analyzed in light of the ratiometric response that the incoherent merger network enables. This could shed light into previously unappreciated functional roles that the above iFFLs cover in their biological context.

IFFLs have been engineered in mammalian cells to achieve constant gene expression despite that transcriptional and translational resources vary in the cell [26] [50]. In particular, it was shown that such an iFFL design can stabilize the expression level of a gene of interest across cell types, thereby possibly enabling constant expression as cells undergo cell type changes, such as in directed differentiation [26]. In the incoherent merger network, global changes in cellular components implicated in gene expression affect the response through an iFFL. This is the first realization of an iFFL that enables robustness of gene expression response to changes in intra-cellular components in bacterial cells. Although our genetic implementation of the incoherent merger network uses a protease as the intermediate species, in other implementations, the intermediate species could be a phosphatase or a scaffold protein that reduces the activity of the output protein.

The incoherent merger network can be widely used in all sensing applications where ratio response is required between any two small molecules for which transcriptional or translational sensors exist. Examples include future smart probiotics that sense stress level and release drugs to improve human performance [14] [15] [16] [12] [13] and programmable organoids where cells need to differentiate into different types while maintaining a desired population size ratio for proper tissue formation [20] [21] [51] [52] [53] [54] [55] [56]. Because in these applications, ratiometric computation operates in orchestration with other genetically encoded components, robustness to changes in cellular resources is critical for proper function. Our incoherent merger network design can therefore be used to engineer multiplexed genetic sense-and-respond systems that are robust to variable cellular context.

## 4 Materials and Methods

### 4.1 Strain and growth medium

Bacterial strain *E. coli* Marionette DH10B (Addgene, #108251) was used to construct and characterize genetic circuits. The growth medium used for construction was LB broth Lennox. The growth medium used for characterization was M9 medium supplemented with 0.4 % glucose, 0.2 % casamino acids, and 1 mM thiamine hydrochloride. Appropriate antibiotics were added according to the selection marker of the plasmid. The final concentration of ampicillin, kanamycin and chloramphenicol are 100, 25, and 12.5 *µ*g*/*mL, respectively.

### 4.2 Genetic circuit construction

The genetic circuit construction was based on Gibson assembly [57]. DNA fragments to be assembled were amplified by PCR using Phusion High-Fidelity PCR Master Mix with GC Buffer (NEB, M0532S), purified with gel electrophoresis and Zymoclean Gel DNA Recovery Kit (Zymo Research, D4002), quantified with the nanophotometer (Implen, P330), and assembled with Gibson assembly protocol using NEBuilder HiFi DNA Assembly Master Mix (NEB, E2621S). Assembled DNA was transformed into competent cells prepared by the CCMB80 buffer (TekNova, C3132). Plasmid DNA was prepared by the plasmid miniprep kit (Zymo Research, D4015). The list of primers and constructs is in Supplementary Figure 1 - Supplementary Figure 3 and Supplementary Table 1.

### 4.3 Microplate photometer

Overnight culture was prepared by inoculating a 80^◦^*C* glycerol stock in 1000 *µL* M9 (+kanamycin) media in a 1.5 ml microcentrifuge tube and grown at 30^◦^*C*, 220 rpm in a horizontal orbiting shaker for 12h. Overnight culture was first diluted to an initial optical density at 600 nm (OD_600nm_) of 0.001 in 200 µL growth medium per well in a 96-well plate (Falcon, 351172) and grown for 1.5 h to ensure exponential growth before induction. The 96-well plate was incubated at 30^◦^*C* in Tecan infinite M Nano+ microplate reader in static condition and was shaken at the “fast” speed for 3 s right before taking OD and fluorescence measurements. The sampling interval was 5 min. Excitation and emission wavelengths to monitor GFP fluorescence were 488 nm and 548 nm, respectively. To ensure enough time to reach steady state GFP/OD signal while the cells were in exponential growth, the cell culture was diluted with fresh growth medium to OD_600nm_ of 0.005 when OD_600nm_ approached 0.135 at the end of each batch. Three batches were conducted for a total experiment time of up to 10 hours until GFP/OD reached steady state. The steady state GFP/OD value was computed from the second batch of each experiment by using the data point with an OD value most close to 0.054 for Figs. 3(a) (g) and Fig. 5(a) - incoherent merger network, and 0.134 for Fig. 4(a) and Fig. 5(a) - broken merging. For the effectors, we choose [IPTG] from 50 to 400 *µ*M and for [Sal] from 0 to 50 *µ*M, which are in the linear range of input/output relationship.

### 4.4 Mathematical modeling and data fitting

Full models of the ODEs are detailed in Supplementary Note 1.1. Data fitting for Figs. 3-5 are performed using MATLAB R2021a (The MathWotks, Inc., Natick, MA, USA) lsqcurvefit function.

### 4.5 Reporting summary

Further information on experimental design is available in the Nature Research Reporting Summary linked to this article.

## 4.6 Data availability

Fluorescence data generated or analyzed during this study are included in the paper and its Supplementary Information files. Essential DNA sequences are provided in Supplementary Figure 1. Full DNA sequences are available on Addgene. A reporting summary for this Article is available as a Supplementary Information file. The source data underlying Figs. 3-5 are provided as a Source Data file. Any other relevant data are available from the authors upon reasonable request.

## 5 Acknowledgements

Funding for this research was provided by the Air Force Office of Scientific Research (AFOSR) under award FA9550-20-1-0044. The views expressed are those of the authors and do not reflect the official guidance or position of the United States Government, the Department of Defense or of the United States Air Force.

